# Polyimidazolium protects against an invasive clinical isolate of *Salmonella* Typhimurium

**DOI:** 10.1101/2022.05.05.490854

**Authors:** Khin K. Z. Mon, Zhangyong Si, Mary B. Chan-Park, Linda J. Kenney

## Abstract

Frequent outbreaks of *Salmonella* Typhimurium infection in both the animal and human population with potential for zoonotic transmission pose a significant threat to the public health sector. The rapid emergence and spread of more invasive multidrug-resistant clinical isolates of *Salmonella* further highlight the need for the development of new drugs with effective broad-spectrum bactericidal activities. Synthesis and evaluation of main-chain cationic polyimidazolium 1 (PIM1) against several gram-positive and gram-negative bacteria have previously demonstrated the efficacy profile of PIM1. The present study focuses on antibacterial and anti-biofilm activities of PIM1 against *Salmonella* both *in vitro* and *in ovo* setting. *In vitro*, PIM1 exhibited bactericidal activity against all tested three strains of *Salmonella* at a low dosage of 8 μg/ml. Anti-biofilm activity of PIM1 was evident with complete inhibition for the initial attachment of biofilms at 16 μg/ml and degradation of pre-formed biofilms in a dose-dependent manner. During the host cell infection process, PIM1 reduces extracellular bacterial adhesion and invasion rates to limit the establishment of infection. Once intracellular, the drug-resistant strain was tolerant and protected from PIM1 treatment. In a chicken egg infection model, PIM1 exhibited therapeutic activity for both *Salmonella* strains with stationary-phase and exponential-phase inocula. Moreover, PIM1 showed a remarkable efficacy against the stationary phase inocula of drug-resistant *Salmonella* by eliminating the bacteria burden in >50% of infected chicken egg embryos. Collectively, PIM1 has demonstrated its potential as a drug candidate for treatment of Salmonella infections, as well as a solution to tackle egg contamination issues on poultry farms.

## INTRODUCTION

*Salmonella enterica* serovar Typhimurium (ST) is a Gram-negative bacterium that can cause disease in a broad range of hosts (1). In humans, ST infection can result in both limiting gastrointestinal disease, as well as extraintestinal systemic infection in immune-compromised patients (2–4). Since ST as a pathogen has high zoonotic potential, with human transmission most commonly occurring through a contaminated food-chain (5), it raises a major public health concern, as well as in the animal and food industries worldwide. The emergence of new, multidrug-resistance (MDR) ST strains (6, 7), further complicates and limits the current antimicrobial therapies available to eradicate this pathogen. *Salmonella* spp. were listed by the World Health Organization (WHO) as among the high-priority antibiotic-resistant bacteria that are in urgent need of new antibiotic development (8, 9). *Salmonella* can exist as either planktonic or as a multicellular aggregate form through biofilm production, which allows prolonged persistence and survival in the host (10). The biofilm state of *Salmonella* is typically antibiotic-tolerant (11, 12), resulting in a chronic bacterial infection that further enhances both the virulence and the transmission rate of the pathogen.

Poultry products are among the highest consumed animal-source products worldwide (13, 14). Consumption of contaminated eggs produced by *Salmonella*-infected layer hens results in foodborne illness, hospitalization, and outbreaks in humans. Therefore, the prevalence of *Salmonella* in the poultry industry has not only a significant economic impact, but it also poses a major threat to public health. To combat against potential pathogens, as well as to promote poultry growth, antibiotics are commonly incorporated into animal food production (14). However, the excessive use or misuse of antibiotic regimes in poultry production can lead to a disastrous outcome of introduction and spread of multi-drug resistance bacteria. Therefore, the development of new therapeutic agents to tackle the increasing number of antibiotic-resistant ST isolates, as well as to prevent bacterial contamination in the food industry is highly desired.

Exploration into the development of synthetic antimicrobial peptides (AMPs) as alternative therapies to existing antibiotics has gained considerable attention over the years, with some promising results. The number of FDA-approved antimicrobial peptides has been steadily increasing over the past decade (15). Common design principles for AMPs consist of cationic hydrophilic-hydrophobic macromolecules, which enables effective targeting and interaction with the negatively-charged bacterial cell surface to promote a contact-based killing mechanism (16). Binding of AMPs to the microbial envelope leads to increased membrane permeability, structural changes (distortion-disruption), leakage of cytoplasmic constituents and eventual bacterial cell death (16–18). Despite its potential for therapeutic usage, there are some limitations associated with the usage of AMPs, including: mammalian cell cytotoxicity, discrepancies between *in vitro* vs *in vivo* efficacy, challenges and potency in human administration, difficulty in synthesis, and the high cost of production (19, 20). An alternate antimicrobial compound known as polyimidazolium (PIM) salt has previously demonstrated a broad spectrum potent antimicrobial activity against several gram-positive and gram-negative bacteria *in vitro* as well as in a murine infection model (21). In the current study, PIM1 was selected for assessment of bactericidal activity against clinical isolates of *Salmonella* Typhimurium including a clinical variant of ST isolated from Vietnam that had acquired a large multidrug-resistance plasmid (6). We also employed an agriculturally relevant model, the chick embryo, to further explore the potential usage of PIM1 in the poultry industry for disease prevention and treatment against the prevalence of *Salmonella*. Overall, PIM1 demonstrated bactericidal activity against clinical isolates of ST under *in vitro* conditions, as well as in a eukaryotic cell infection assay and an *in ovo* infection model.

## RESULTS

### Antibiotic drug panel screening of Salmonella *Typhimurium* strains

A preliminary screening of ST strains against antibiotic drug panels demonstrated high scores for the MBC assay and half-maximal inhibitory concentration (IC_50_) value, most notably for the MDR strain (Tables 1A-B). Out of twelve antibiotics screened, eight drugs scored an IC_50_ value higher than 50 µg/ml, attributing to its acquisition of a large multidrug-resistance plasmid (6). The loss of the MDR plasmid in the plasmid-cured strain ΔMDR, drastically improved the sensitivity and reduced the score of the MBC assay with antibiotics (Table 1B).

**Table 1:** (A) IC_50_ value of *Salmonella* strains screened for 12 antibiotics. (B) MBC assay scores of *Salmonella* strains screened for the same 12 antibiotics.

| Antibiotics | IC<sub>50</sub> (µg/ml) |  |  |
| --- | --- | --- | --- |
|  | 14028 | MDR | ΔMDR |
| Ampicillin | 3.498 | 4026 | 5.131 |
| Kanamycin | 1.473 | 29.07 | 2.378 |
| Streptomycin | 6.304 | 5.338 | 5.356 |
| Carbenicillin | 0.6905 | 167.1 | 1.499 |
| Gentamycin | 0.03811 | 16.54 | 3.648 |
| Chloramphenicol | 1.845 | 136.8 | 5.105 |
| Tetracyclin | 1.001 | 58.88 | 61.73 |
| Trimethoprim | 4.583 | 494.4 | 14.77 |
| Novobiocin | 133.9 | 152.4 | 94.73 |
| Spectinomycin | 33.35 | 89.01 | 40.42 |
| Erythromycin | 21.87 | 26.81 | 12.52 |
| Nalidixic acid | 3.794 | 350.6 | 247.2 |

| Antibiotics | MBC (µg/ml) |  |  |
| --- | --- | --- | --- |
|  | 14028 | MDR | ΔMDR |
| Ampicillin | 25 | 6400 | 50 |
| Kanamycin | 6.25 | 800 | 6.25 |
| Streptomycin | 25 | 50 | 25 |
| Carbenicillin | 6.25 | 400 | 6.25 |
| Gentamycin | 3.125 | 200 | 1.56 |
| Chloramphenicol | 50 | 1600 | 100 |
| Tetracyclin | 25 | 100 | 100 |
| Trimethoprim | 100 | 3200 | 1600 |
| Novobiocin | 1600 | 800 | 1600 |
| Spectinomycin | 6400 | 102400 | 6400 |
| Erythromycin | 800 | 800 | 1600 |
| Nalidixic acid | 12.5 | 6400 | 6400 |

### In vitro bactericidal activity of PIM1

The antimicrobial activity of PIM1 was evaluated against ST lab strain 14028s, the multidrug-resistant (MDR) clinical isolate 20081, and the plasmid-cured strain of 20081, resulting in the loss of plasmid-encoded antibiotic resistance genes (ΔMDR) *in vitro*. A low minimum inhibitory concentration (MIC) value of 4 µg/ml demonstrated the killing efficacy of the PIM-1 polymer against all three *Salmonella* strains (Figure 1A). The corresponding minimum bactericidal concentration (MBC) values where there was no viable bacterial growth detected on the plates was 8 µg/ml for all three strains (Figure 1A). Therefore, PIM1 displayed potent antimicrobial activity against *Salmonella* Typhimurium isolates at low doses of 4-8 µg/ml. To further evaluate the killing kinetics of PIM1, all three strains were grown in the presence of PIM1 at 2x and 4x the MBC value obtained (16 µg/ml and 32 µg/ml). All three *Salmonella* strains were rapidly killed within 15 minutes at 32 µg/ml, while it took 30 minutes at 16 µg/ml (Figure 1B-D). A steady increase in bacterial growth over the same time period was observed in the non-PIM1 treated group, confirming the bacterial killing effect of PIM1 in the treated groups.

**Figure 1:**
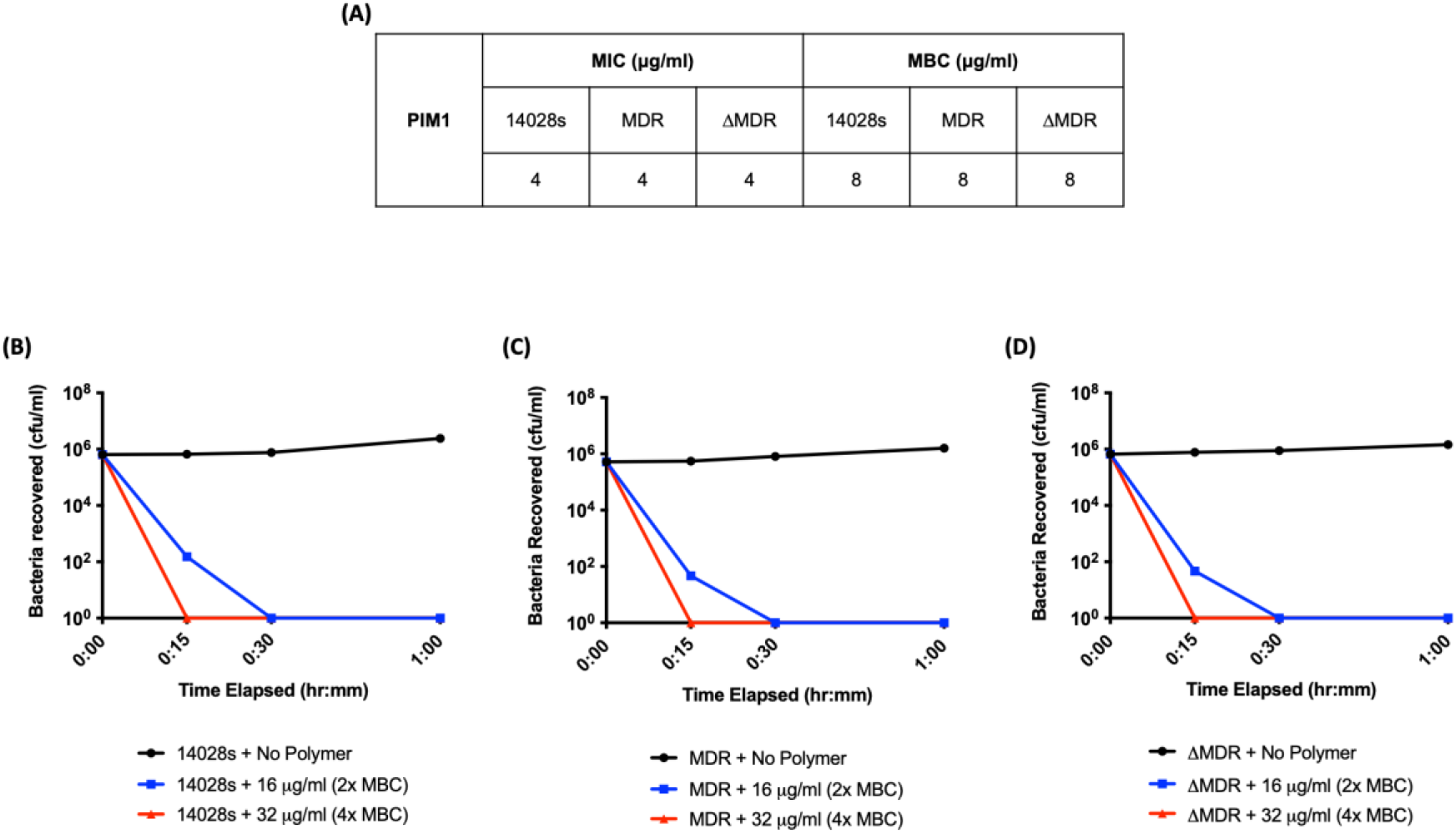
Bactericidal efficiency of PIM-1 against three ST isolates. (A) Bacterial cultures in Muller-Hinton Broth were treated overnight with a PIM1 dose at 2x increments and the observed Minimal Inhibitory Concentration (MIC) and Minimal Bactericidal Concentration (MBC) for all three strains was recorded. The data are representative of four replicates. (B-D) Bacterial killing kinetics of PIM1 was measured at 2x and 4x concentration of the MBC value (16 µg/ml and 32 µg/ml) at 15 min, 30 min and 1h intervals compared to the bacterial growth rate (untreated). The data plotted were the mean of triplicate plate counts of the number of viable bacteria recovered at each time interval.

### Anti-biofilm activity of PIM1

A PIM1 inhibitory effect on *Salmonella* biofilm formation, as well as degradation, was quantified via a crystal violet assay. In the absence of PIM1, all tested bacteria were able to form biofilms under biofilm growth conditions. Specifically, MDR and ΔMDR strains were observed to have enhanced biofilm thickness measured in units of crystal violet absorbance compared to the lab strain 14028s (Figure 2A-B). A PIM1 concentration of as low as 16 µg/ml significantly inhibited biofilm formation (Figure 2A, p<0.05 for 14028s and p<0.0001 for MDR and ΔMDR). Since MDR and ΔMDR strains were observed to be a hyper-biofilm variant, the ability of PIM1 to effectively degrade and eradicate pre-established biofilms was also explored. A dose-dependent effect of PIM1 on biofilm degradation was observed, with a maximal reduction in biofilms for all tested bacteria at 1024 µg/ml (Figure 2B, p<0.0001). PIM1 effects on biofilm formation and degradation were further confirmed by fluorescence microscopy imaging with SYTO-9 staining (nucleic acid stain). The z axis measurements from the biofilm image stacks validated that the biofilm thickness of MDR and ΔMDR strains were indeed much thicker (∼two-fold) than the biofilm of 14028s. A reduction in biofilm thickness for all tested strains was also evident from the z-axis scale, with post-PIM1 treatment at 1024 µg/ml (Figure 2C). All three bacterial strains showed a statistically significant decrease in overall biofilm volume post-PIM1 treatment (Figure 2D-F, p<0.001 for 14028s & ΔMDR and p<0.05 for MDR).

**Figure 2:**
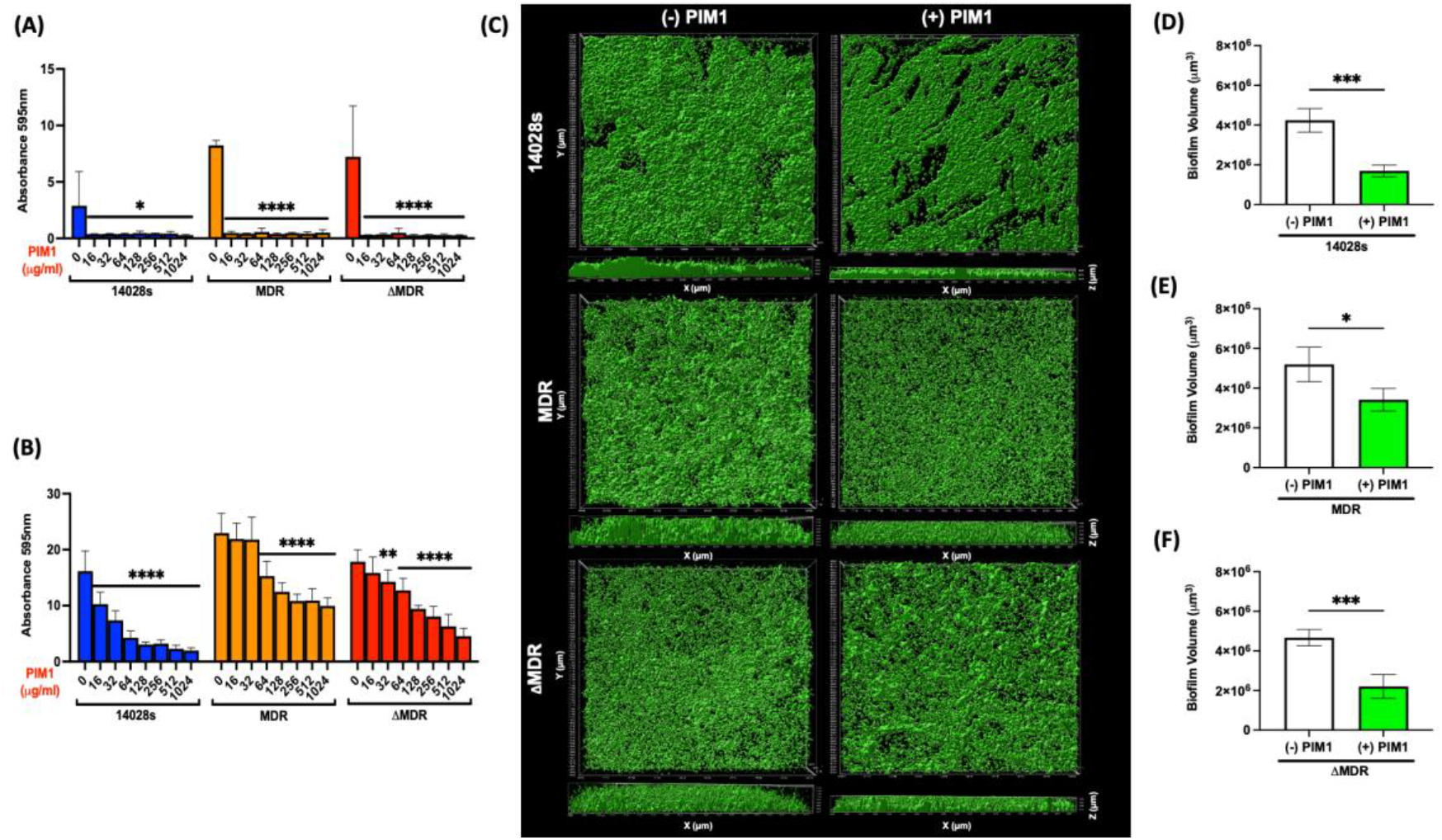
Anti-biofilm activity of PIM1 against three ST strains. (A) PIM1 inhibitory effect on biofilm formation was measured using crystal violet staining for all three strains (14028s, MDR and ΔMDR) at various concentrations of PIM1. The results represent the mean ± SD, *P-value<0.05 between untreated and PIM1-treated groups for 14028s, ****P-value<0.0001 between untreated and PIM1-treated groups for MDR and ΔMDR strains (Dunnett’s multiple comparisons test). (B) Dose-dependent effects of PIM1 on biofilm degradation as measured by crystal violet staining. The results represent the mean ± SD, ****P-value<0.0001 between untreated and all PIM1-treated groups for 14028s, ****P-value<0.0001 between untreated and PIM1-treated groups for MDR with the exception of 16 μg/ml and 32 μg/ml PIM1-treated groups (not significant), ****P-value<0.0001 between untreated and PIM1-treated groups for ΔMDR strains, except for 16 μg/ml (not significant) and 32 μg/ml (**P-value<0.01) (Dunnett’s multiple comparisons test). (C) Three-dimensional (3D) modeling of *Salmonella* biofilms stained with SYTO-9 (green) in the absence (untreated PIM1) or presence of PIM1 at 1024 μg/ml using the surface function in Imaris software with xy-plane and xz-plane views (D-F) The biofilm volume for the post-PIM1 treated group (1024 μg/ml) for all three strains was significantly reduced in comparison to the untreated group. The results represent the mean ± SD, ***P-value<0.001 between the untreated group and the post-PIM1 treated group for 14028s and ΔMDR strains, *P-value<0.05 for the untreated group and the post-PIM1 treated group for MDR (t test).

### PIM1 interferes with adhesion and invasion, but not intracellular replication of the invasive clinical isolate strain MDR

The cytotoxicity of PIM1 on mammalian cells was first assessed with HeLa cells. At the highest concentration of PIM1 (1024 µg/ml), HeLa cell survival was drastically reduced, with detection of ∼1% viable cells. At lower PIM1 concentrations, the HeLa cell survival rate remain largely unaffected by the PIM1 treatment, with viability of 93% (MIC), and 83% (MBC), respectively (Figure 3A). The half-maximal inhibitory concentration of the drug, also known as the IC_50_ value, at which a 50% reduction in HeLa cell viability occurs was estimated to be 130 µg/ml. Hence, we used PIM1 concentrations (MIC & MBC) that were determined not to have a high cytotoxicity effect on HeLa cells for the subsequent *Salmonella* cell infection assay. To determine the ability of PIM1 to directly interfere with the ability of *Salmonella* to attach and invade cells during infection, PIM1 was added at the same time as the bacterial inoculation. PIM1 exhibited a strong inhibitory activity against cell adhesion of both 14028s and MDR strains (p<0.0001) at all tested concentrations compared to the untreated control group (Figure 3B, D). PIM1 effectively blocked bacterial adherence to HeLa cells at a MIC treatment level (4 µg/ml) by significantly reducing the total number of attached bacteria by ∼20,000 fold for both strains (14028s and MDR). For *Salmonella* invasion and replication assessment, a gentamicin protection assay was performed to obtain the intracellular bacterial count. Since MDR is a gentamicin-resistant strain, it was replaced with the plasmid-cured MDR strain (ΔMDR) that has lost the gentamicin-resistance phenotype for this assay. A steady decrease in the *Salmonella* cell invasion rate was observed in a PIM1 dose-dependent manner for 14028s and ΔMDR (p<0.0001) compared to untreated *Salmonella-*infected HeLa cells (Figure 3B, D). To assess the ability of PIM1 to target and eliminate the intracellular bacteria during an active infection, PIM1 was added later to *Salmonella-*infected HeLa cells, after the gentamicin protection step. At 2h post-infection, a higher treatment dose (MBC = 8 µg/ml) allowed PIM1 to enter host cells, which led to a reduction in the intracellular bacterial load of *Salmonella* 14028s (Figure 3B, p<0.01). On the other hand, the intracellular replication rate of *Salmonella* ΔMDR within HeLa cells remained unaffected, even after the PIM1 treatment dose was incrementally increased (Figure 3D). Taken together, these results indicated that although PIM1 can effectively inhibit the host cell attachment and invasion of bacteria, it is largely unable to target and eradicate intracellular bacteria. PIM1 interference with *Salmonella* infection stages in host cells was further confirmed via confocal fluorescence microscopy. A PIM1-fluorescein isothiocyanate (FITC) conjugate was used in *Salmonella* infections of HeLa cells to visualize and further validate the PIM1 inhibitory activity during an active infection. Microscopy images corroborated the quantitative CFU dataset, where PIM1 treatment was largely effective in visibly reducing the bacterial load of both strains for adhesion and invasion stages in a dose-dependent manner (Figure 3C, E).

**Figure 3:**
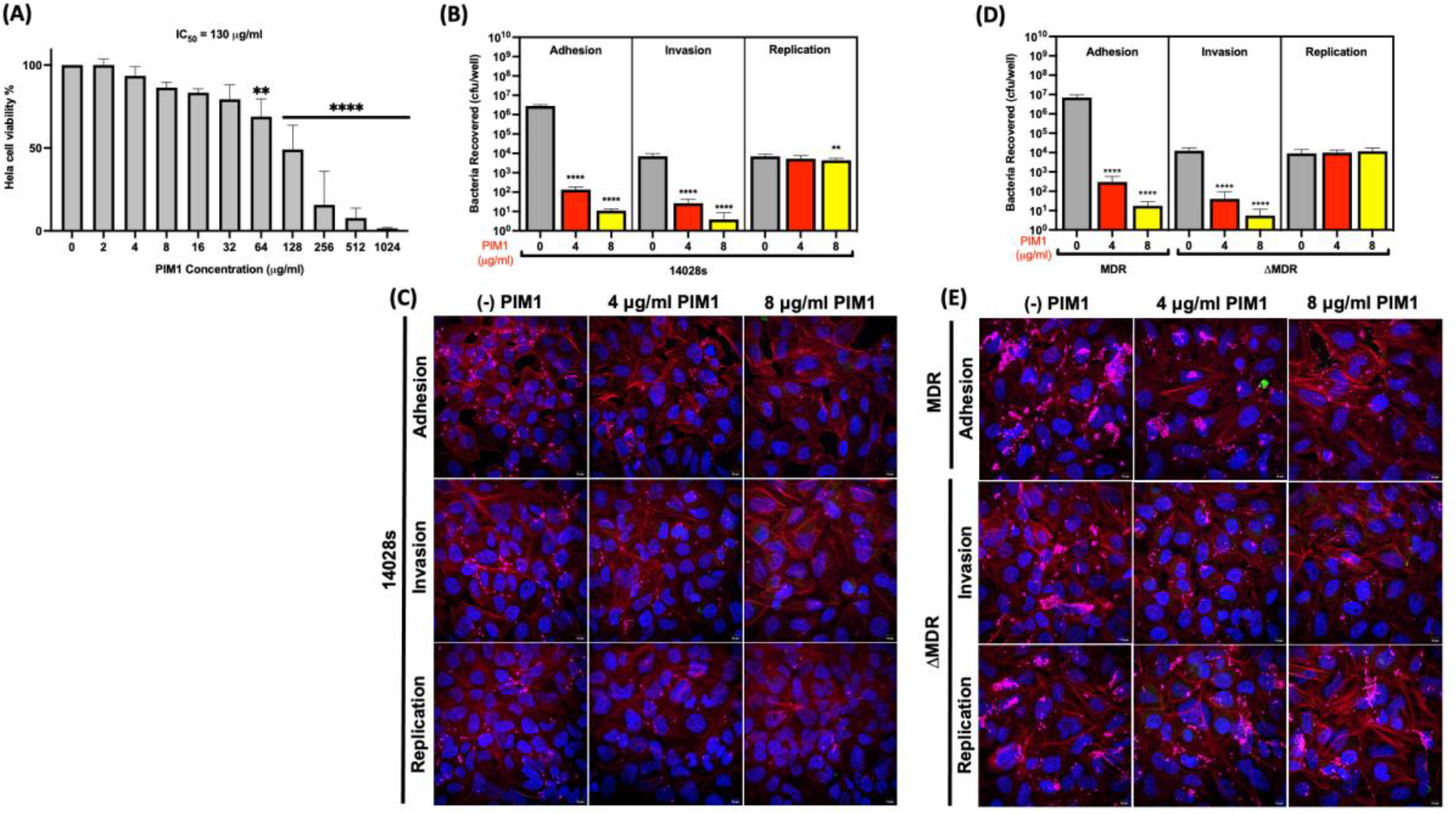
The effect of PIM1 on mammalian cell cytotoxicity and its inhibitory activity against *Salmonella* during cell infection. (A) IC_50_ cytotoxicity value of HeLa cells treated with PIM1 was determined using the MTT assay. The results represent the mean ± SD (n = 3), **P-value<0.01 between untreated and PIM1 treated HeLa cells (64 μg/ml), ****P-value<0.0001 between untreated and PIM1 treated HeLa cells at 128, 256, 512 & 1024 μg/ml. (B) Recovery of *Salmonella* 14028s during adhesion, invasion, and intracellular replication in the presence or absence of PIM1 at MIC and MBC concentrations. The results represent the mean ± SD (n = 2), **P-value<0.01, ***P-value<0.001 and ****P-value<0.0001 between untreated (-) and PIM1-FITC treated groups (4 and 8 μg/ml) (Dunnett’s multiple comparisons test). (C) Representative confocal microscopy z stack images of HeLa cells (stained with DAPI = blue and Phalloadin 568 = red) infected with *Salmonella* 14028s (LPS 647 = magenta) in the absence or presence of PIM1-FITC (green) at MIC and MBC concentrations during different stages of infection. (D) CFU assay of *Salmonella* MDR 20081 adhesion, invasion, and replication rate (ΔMDR tested in place of MDR for the gentamicin protection assay) in the presence or absence of PIM1 at MIC, and MBC concentrations. The results represent the mean ± SD (n = 2,), **P-value<0.01, ***P-value<0.001 and ****P-value<0.0001 between untreated (-) and PIM1-FITC treated groups (4 and 8 μg/ml) (Dunnett’s multiple comparisons test). (E) Representative confocal microscopy z stack images of HeLa cells (stained with DAPI = blue and Phalloidin 568 = red) infected with *Salmonella* MDR and ΔMDR (LPS 647 = magenta) in the absence or presence of PIM1-FITC (green) at MIC and MBC concentrations during different stages of infection.

### PIM1 demonstrated therapeutic activity in chicken egg model of Salmonella infection

The ability of PIM1 to control and reduce *Salmonella* colonization in the chick embryo chorioallantoic membrane (CAM) model at 16 µg/egg dose (2x MBC) was assessed using both stationary phase and exponential phase bacterial inocula. The CAM model was comprised of multiple steps and was performed over 12 days from egg incubation to tissue harvesting, as illustrated in Figure 4A. *In ovo* imaging of CAMs infected with mCherry-expressing *Salmonella* 14028s confirmed the signal detection of red fluorescence at the bacterial inoculation site on filter papers. In PIM1-FITC treated CAMs, the red fluorescence bacteria signal was eliminated and replaced by a PIM1 green fluorescence signal at 1dpi compared to the untreated CAM, for both growth-phase inocula (Figure 4B). Fluorescence *in ovo* images corroborated the significant decrease in bacterial load in *Salmonella* 14028s-infected CAMs. With either inocula, there was a significant reduction in the total bacterial burden recovered from PIM1-treated 14028s CAMs compared to untreated 14028s CAMs (Figure 4C-D, stationary-phase inoculum = p<0.0001, exponential-phase inoculum = p<0.0001). Furthermore, complete bacterial clearance in both the CAM and the liver was detected in three out of ten chick embryos for 14028s exponential phase inocula post-PIM1 treatment (Figure 4D). PIM1 also exhibited a potent activity against the MDR strain *in ovo* with a significant decrease in the bacterial load for MDR-infected CAMs (Figure 4E-F, stationary-phase inoculum = p<0.0001, exponential-phase inoculum = p<0.001). Notably, a stationary phase inocula of MDR was more susceptible to the PIM1 bactericidal effect than an exponential phase inocula. Complete bacterial clearance was detected in four out of seven (>50%) infected chick embryos (both CAM and liver) (Figure 4E). Preliminary *Salmonella* infection experiments in the immunodeficient CAM model have indicated that *Salmonella* was capable of rapidly disseminating to internal organ sites of the chick embryo within1 hpi. Thus, the ability of PIM1 to limit or reduce the disseminated bacterial load in the liver was also evaluated. A significant reduction of bacteria in the liver was detected, with complete bacterial clearance in three out of ten and four out of eight livers for 14028s and MDR strains, respectively (exponential phase inoculum, Figure 4D and 4F, p<0.0001). For the stationary phase inoculum, PIM1 treatment exhibited greater efficacy in MDR than 14028s-infected chick embryos by completely clearing the bacterial burden in four out of seven livers (Figure 4E, p<0.001). Although complete bacterial clearance in the livers of 14028s-infected chick embryos was not detected, there was a significant reduction in the total bacteria burden in the liver post PIM1 treatment (Figure 4C, p<0.05).

**Figure 4:**
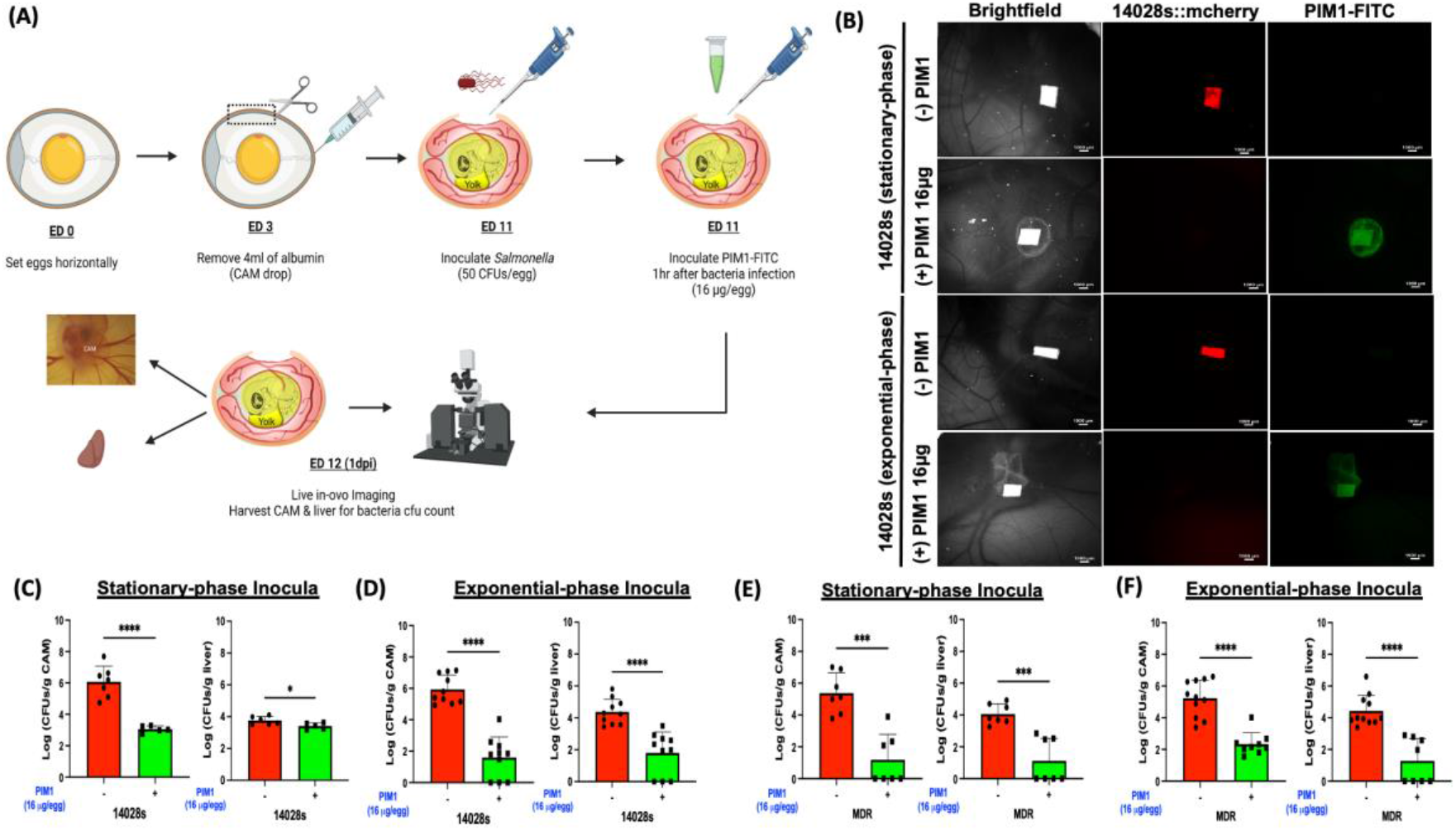
PIM1 reduces *Salmonella* infection in a chicken egg embryo model. (A) The experimental outline of the chick embryo chorioallantoic membrane (CAM) infection model. (B) Representative live fluorescence stereomicroscope images *in ovo* with *Salmonella* 14028s::mCherry (red) with or without PIM1-FITC (green) 16 μg/egg at 1 dpi with stationary phase or exponential phase inocula. (C-F) Quantification of bacteria recovered from the CAMs and livers of chick embryos infected with stationary phase or exponential phase inocula of 14028s and MDR in the absence or presence of PIM1 treatment at 16 μg/egg, 1dpi. The results are presented as the exponential of the CFUs, mean ± SD, *P-value<0.05, ***P-value<0.001 and ****P-value<0.0001 between untreated (-) and PIM1-FITC treated groups (16 μg/egg) (t test).

## DISCUSSION

The emergence and high prevalence of multidrug-resistant bacteria are a global public health threat, as it decreases the effectiveness of conventional antibiotic treatment for pathogen infections. The widespread usage of antibiotics has also led to the natural selection process of microorganisms acquiring multi-drug resistance traits or genes, which further complicates the available treatment options (22–24). The development of imidazolium salts as an alternative antimicrobial compound has generated promising results and advances in recent years (25–27). It offers a solution for combating multidrug-resistant bacterial pathogens and bacterial-associated infections. We have previously reported on the potency of a series of main-chain cationic polyimidazolium (PIMs) against several gram-positive and gram-negative bacteria (21). In the current study, we demonstrated the efficacy of a selected PIM series, PIM1, against invasive clinical isolates of *Salmonella* Typhimurium 14028s, MDR and its plasmid-cured counterpart ΔMDR.

Under *in vitro* testing conditions with PIM1, bactericidal activity against a planktonic population of *Salmonella* strains was observed with low MIC and MBC scores of 4-8 µg/ml. This was identical to the MIC and MBC scores with PIM1 with *Escherichia coli*, another gram-negative food-borne bacterial pathogen (21). The high sensitivity of the MDR strain towards PIM1 with a low MBC value, as well as rapid killing kinetics demonstrated the efficacy of the compound (Figure 1A, C).

The lifestyle switch from planktonic to the biofilm state allows pathogens such as *Salmonella* to survive under harsh environmental conditions and promote chronic carriage, as well as transmission of disease (28, 29). Biofilms are a multicellular community that are embedded in a self-produced matrix of extracellular polymeric substances (EPS), growing on either biotic or abiotic surfaces that are difficult to eradicate, with high tolerance towards antimicrobial agents (12, 29, 30). In the current study, the antibiofilm effect of PIM1 was evident, with inhibition of biofilm formation at all tested concentrations (Figure 2A) and a dose-dependent effect on degradation of pre-formed biofilms in vitro for all three strains, as measured by the crystal violet assay (Figure 2B). The MDR strain, and its plasmid-cured counterpart ΔMDR, exhibited a hyper-biofilm forming phenotype compared to the lab strain 14028s. Both MDR strains have much thicker biofilms (visible to the naked eye) that were confirmed by a higher absorbance in the crystal violet assay, as well as in the z axis scale from fluorescence images (Figure 2A-C). In addition, MDR and ΔMDR strains form biofilms at the air-liquid interface, while 14028s biofilms are submerged at the bottom of the well. Biofilms that form at the air-liquid interface have been reported to be much thicker and more resistant to cleaning agents (31, 32). Regardless, the PIM1 effectiveness in degrading pre-formed biofilms was evident in terms of a reduction in thickness, as well as biofilm volume, in dense, three-dimensional (3D) architecture images of all *Salmonella* strains, including the hyperbiofilm variants (Figure 2C-F). However, PIM1 was unable to completely eradicate the biofilm at the highest tested concentration of 1024 µg/ml and may require a much higher concentration of PIM1 or a longer treatment time.

The host infection process by the intracellular pathogen *Salmonella* occurs in different stages: 1) adhesion or association of bacteria with eukaryotic cells, 2) host cell invasion and 3) intracellular replication. These events can be evaluated *in vitro* based on infection time intervals, a gentamicin protection assay and post-gentamicin incubation timeline (33). Therefore, the inhibitory activity of PIM1 on *Salmonella* adhesion, invasion and replication events were evaluated with some modifications. Attachment of the bacterial pathogen to the host cell surface is the essential first step in the pathogenesis of infection, while invasion of bacteria into host cells determines the subsequent bacterial survival and establishment of the infection within the host (34). PIM1 interferes early in the bacterial infection process (adhesion and invasion stages) by exhibiting bactericidal activity towards extracellular *Salmonella* strains at both MIC and MBC levels in a dose-dependent manner. In contrast, intracellular *Salmonella* were largely protected against PIM1 killing during the later stage of the infection process, replication (Figure 3B-E). PIM1 treatment at a higher dosage of 8 µg/ml was effective in significantly reducing the bacterial load of 14028s at the 2h replication stage, indicating the entry of PIM1 across the host cell membrane to target intracellular bacteria. However, a mammalian cytotoxicity assay also indicated that PIM1 triggered higher cell toxicity of 17% at this concentration compared to 7% at an MIC of 4 µg/ml (Figure 3A). It is also worth mentioning that the PIM1 concentration used in *Salmonella* cell infection assays was much lower than the IC_50_ cytotoxicity value of HeLa cells (4 and 8 µg/ml used in infection assay compared to IC_50_ of 130 µg/ml). An approximately sixteen-fold increase in the PIM1 concentration would be required for a 50% reduction in HeLa cell viability. Additionally, four other mammalian cell lines have been previously tested and a much higher IC_50_ concentration (>800 µg/ml) was reported (21). These results suggest that PIM1 has targeted bactericidal activity, while exhibiting minimal cytotoxicity effects towards mammalian cells. Furthermore, the antimicrobial activity of PIM1 at the early stages of the infection process played a crucial role in suppressing bacterial colonization and disrupted the establishment of infection within the host.

*Salmonella-*associated food borne outbreaks are often traced back to contaminated eggs or egg-derived products, which serve as an important food vehicle for transmission of infection (35–37). Positive *in vitro* antimicrobial activity of PIM1 led us to further explore the protective potential of PIM1 administration in a chicken egg model, an agriculturally relevant model for control of *Salmonella* infection in the poultry industry. The efficiency of antimicrobial treatment and its susceptibility is influenced by many factors, including the growth phase of the bacterial inoculum. Stationary-phase bacteria typically exhibit a more resistant phenotype towards antimicrobial agents compared to exponential phase populations. This is because stationary phase bacteria undergo changes at both molecular and cellular levels to withstand harsh environmental conditions (38–40). Hence, PIM1 treatment was tested against infected chicken egg embryos of either stationary phase or exponential phase inocula. PIM1 was bactericidal *in ovo* with a significant reduction in the bacterial load at both CAM and liver sites (Figure 4C-F). Such an extraordinary bactericidal effect of PIM1 was observed after administration of a low dose (16 µg per egg), irrespective of the inoculum. Since the chicken egg embryo is an immunodeficient model (41), the possibility of triggering host immunity to limit bacterial colonization during a short infection time interval is an unlikely scenario. Instead, reduction of the bacterial load in the chick embryo model was most likely a direct consequential effect of PIM1 bactericidal activity.

In summary, PIM1 demonstrated potent efficacy against *Salmonella* Typhimurium wildtype 14028s, a multi-drug resistant clinical variant MDR and its ΔMDR derivative. Specifically, PIM1 was effective at a low concentration towards planktonic populations, while also inhibiting biofilm formation of all strains, including multi-drug resistant hyper-biofilm variants. A biofilm degradation capability was observed that was dose-dependent, with the most drastic degradation observed at the highest concentration tested (1024 µg/ml). PIM1 interfered with bacterial adhesion and invasion events during eukaryotic cell infection, further highlighting its ability to target extracellular bacteria to limit pathogen survival and establishment of infection within the host. The effective antimicrobial activity of PIM1 was consistent in both *in vitro* and *in vivo* settings. In a chicken egg embryo model of *Salmonella* infection, a single low dose of PIM1 treatment effectively decreased the bacterial load in both the CAM and the liver during infection at 1dpi. Although PIM1 did not show any evidence of toxicity towards chick embryos in the current study (at the low administered dose of 16 µg per egg), PIM1 exhibited acute toxicity in mice when administered intraperitoneally at 6 mg/kg dose (21). A modified derivative of PIM1, PIM1D with reduced hydrophobicity compared to PIM1 resolved the toxicity issue in a murine sepsis infection model. Therefore, the better safety profile of PIM1D suggests a promising therapeutic applicability of the agent as an alternative treatment for both human and poultry associated *Salmonellosis*.

## MATERIALS AND METHODS

### Minimum Inhibitory Concentration (MIC) and Minimum Bactericidal Concentration (MBC) of PIM1

Bacterial cultures were prepared in Muller Hinton (MH) broth and incubated overnight at 37°C at 225 rpm. The overnight cultures were resuspended into fresh MH broth (1:100) and incubated at 225 rpm, 37°C to reach the mid-exponential phase of growth. After incubation, bacterial cultures were diluted to ∼5 × 10^5^ cfu/ml. The PIM1 polymer was serially diluted in two-fold increments in MH broth to a concentration ranging from 0 to 64 μg/ml. Mixed suspensions of the polymer and bacteria were incubated overnight at 37°C without shaking in a 24-well microtiter plate. The MIC was determined with no visible growth of bacteria in a spectrophotometer (OD_600_ nm). Bacteria without added polymer provided a positive growth control. For the MBC assay, a spot assay was performed with 10 µl of the bacterial suspension on an LB agar plate in the presence of varying polymer concentrations (determined from the MIC assay) and incubated at 37°C overnight. The MBC was recorded as the lowest concentration of polymer that killed 100% of the bacterial inoculum.

### Killing kinetics of PIM1

Overnight bacterial cultures were resuspended in fresh MH broth (1:100) and incubated at 37°C with shaking at 225 rpm until the cultures reached mid-exponential phase. The bacterial suspension was then diluted to 1 × 10^5^ cfu/ml in MH broth with a polymer concentration of two times or four times the MBC value obtained (16 μg/ml and 32 μg/ml). A bacterial culture without polymer served as a positive growth control. Bacteria and polymer mixtures were incubated at 37°C, 150 rpm for time intervals of 15 min, 30 min and 1h. At each timepoint, 100 µl of the mixture was serially diluted onto LB agar plates. The number of bacteria recovered were counted after overnight incubation at 37°C. Results in triplicate were averaged and plotted.

### Biofilm quantification with crystal violet assay

A crystal violet assay for biofilm quantification in the absence and presence of PIM1 was performed as described previously (28) with some modifications. Briefly, for the biofilm inhibition assay, 2 µl of the overnight culture was inoculated into 198 µl of LB medium without salt with serial two-fold dilution of PIM1 (0, 16, 32, 64, 128, 256, 528, 1024 µg/ml) in a 96-well polystyrene plate and grown statically at 30°C for two days. For the biofilm degradation assay, the growth medium from pre-established biofilms was removed, washed and replaced with fresh medium containing two-fold serial dilution of PIM1 for further 24h incubation at 30°C. Planktonic cells were then removed and the attached biofilms in each well were stained with crystal violet. The crystal violet solution was then aspirated, and wells were washed one time with PBS followed by the addition of absolute ethanol to solubilize the crystal violet. Ten-fold dilutions were measured for absorbance at 595 nm using a Biorad plate reader. The experiment was performed twice with quadruplicates for all three strains under each treatment condition.

### Fluorescence imaging of biofilms

Biofilms were grown in the µ-slide 18 wells, glass bottom coverslip (Catalog No. NC1752257, Ibidi) for two days at 30°C. Planktonic cells were removed from the well and replaced with fresh growth medium containing 1024 µg/ml of PIM1. Fresh growth medium only was added into control wells for the non-treatment group. The µ-slide 18 wells were further incubated for 24h of PIM1 treatment at 30°C. Each well was then washed and stained with 2 µM SYTO-9 green for a minimum of 30 minutes before fluorescence images were acquired. Image acquisition was performed using an Olympus Super Res Spinning Disk, SpinSR-10 microscope at 480/500 nm (40x magnification). Image stacks were acquired from seven representative areas of the biofilm surface from each well. The experiment was repeated twice in duplicate for all three strains in the presence or absence of PIM1. Image stacks were reconstructed three dimensionally and biofilm volume analysis was performed using the surface function of Imaris software 9.7.

### Mammalian cell cytotoxicity assay

The PIM1 cytotoxicity on HeLa cells was determined using a 3-[4,5-dimethylthiazol-2-yl]-2,5 diphenyl tetrazolium bromide (MTT) assay. Briefly, cells were first seeded in 94-well cell-culture plates at a density of 2 × 10^4^ cells per well and incubated at 37°C with 5% CO_2_ for 24h. The cell culture medium was replaced with fresh medium in the presence of PIM1 (at 2, 4, 8, 16, 32, 64, 128, 256, 528, 1024 µg/ml) and incubated for a further 24h. Untreated cells (without PIM1) served as a negative control, while empty wells with cell medium alone served as a blank control. The next day, the cell medium was aspirated and replaced with fresh medium containing MTT and incubated for 4 h. The MTT solution was then carefully aspirated and DMSO was added to solubilize. The absorbance readout at 595nm was recorded using a Biorad plate reader. The percentage of viable cells was calculated using the ratio of the average OD_595_ of treated cells to the average OD_595_ of untreated cells.

### Adhesion, invasion and intracellular survival & replication assays

The ability of PIM1 to interfere with *Salmonella* attachment, invasion as well as intracellular replication within HeLa cells during the infection process was investigated with PIM concentrations of MIC (4 μg/ml), and MBC (8 μg/ml). Briefly, 24-well plates were seeded with 5 × 10^4^ cells/well. Subcultures of *Salmonella* were added to the wells in Dulbecco’s modified Eagle’s medium (DMEM). The infection assay was carried out as previously described (42) with some modifications. For the adhesion and invasion assay, PIM1 was added to the wells at the same time as *Salmonella* and incubated for 30 min after being centrifuged at 500 x g for 10 min. The infection process was stopped for adhesion analysis by lysing the cells with 0.1% Triton X-100 for 10 min, while invasion analysis was further treated with gentamicin (100 μg/ml) for 1h to kill the extracellular bacteria before cells were lysed for plating. Since the MDR strain is resistant to gentamicin, a ΔMDR strain in which the multi-drug resistant plasmid was cured from the parent strain (Y. Gao & L.J. Kenney, unpublished results) replaced the MDR strain for both invasion and replication assays where gentamicin treatment was involved. The effect of PIM1 on targeting intracellular bacteria was determined by the addition of PIM1 to the wells after the initial high concentration of gentamicin (100 μg/ml) treatment for replication analysis. A lower concentration of gentamicin (20 μg/ml) was added to the untreated group to eliminate extracellular bacterial growth. Infected cells were washed twice with PBS and lysed with 0.1% Triton X-100 for 10 min at 30 min post-infection (adhesion), 1h post-gentamicin treatment (invasion), and 2h (replication). Lysates were serially diluted and plated on LB agar to obtain the colony-forming units (CFU).

### Immunostaining

Immunostaining was carried out on fixed adherent HeLa cells on coverslips at different stages of infection both in the absence and presence of PIM1-FITC treatment. Fixed HeLa cells were incubated with primary antibody solution in PBS with 5% bovine serum albumin and 0.1% saponin. *Salmonella* strains were stained with monoclonal rabbit anti-LPS (1:500 dilution). After 1h, primary antibody solution was removed, and cells were gently washed three times with PBS containing 0.1% Tween-20. Secondary antibody solution, which included donkey anti-rabbit 647 (Invitrogen, A-31573), Alexa Fluor phalloidin 568 (Invitrogen A-12380) and DAPI (Invitrogen, H3570) in PBS-BSA-saponin buffer was incubated for 1h at room temperature in the dark. After subsequent wash steps, coverslips containing the adherent stained HeLa cells were inverted onto a microscope slide containing ProLong gold antifade reagent (Invitrogen, P36934), sealed with nail polish, and allowed to dry overnight in the dark before image acquisition with an Olympus Super Res Spinning Disk, SpinSR-10 microscope at 100x magnification.

### Chicken egg embryo infection model

Specific pathogen free (SPF) embryonated chicken eggs were purchased from Charles River Laboratories. The chick chorioallantoic membrane (CAM) drops on embryonic development day 3 (ED3) by removing 4 ml of albumin using a thin needle inserted at the apex of the egg. A small window was made on the top of the eggshell. On ED10, a bacterial inoculum was applied directly on a small piece of sterile filter paper that was placed on top of the CAM. For a stationary phase inoculum, an overnight bacterial culture was prepared, while a subculture was used for the preparation of the exponential-phase inoculum. PIM1 (16 μg) was then applied on the same spot as the bacterial inoculum deposition site at 1h post-infection. At one day post-infection (1 dpi), the CAM layer and the embryo liver were harvested. Samples were then homogenized and serially diluted to determine bacterial numbers by the colony counting method. Imaging of live embryos that were infected with mCherry-tagged ST 14028s was also performed at 1 dpi to detect the fluorescence signal of bacteria on filter papers with or without PIM1-FITC treatment using a stereomicroscope.

## ACKNOWLEDGMENTS

We thank Dr. Stuti Desai for her expertise and guidance with the biofilm aspects of this study and Dr. E.P. Greenberg (University of Washington) for his comments on the manuuscript. Early studies were supported by a Research Centre of Excellence grant from the Ministry of Education to the Mechanobiology Institute at the National University of Singapore. KKZM was supported by CPRT RP200650 to LJK. Z.S. and M.B.C. were funded and supported by the Singapore MOE Tier 3 grant (MOE2018-T3-1-003).

## Abbreviations

h: hour
min: minutes
DMEM: Dulbecco’s modified Eagle’s medium
ED: embryonic development day
dpi: day post-infection
SD: standard deviation
CAM: chorioallantoic membrane

